# Lesion-Derived Estimated Functional and Structural Connectomes Predict Post-Stroke Recovery Outcomes

**DOI:** 10.64898/2026.10.30.762945

**Authors:** Christie Gillies, Keith Jamison, Aaron D. Boes, Amy Kuceyeski

**Affiliations:** Department of Radiology, Weill Cornell Medicine, New York, NY, USA; Computational Biology, Cornell University, Ithaca, NY, USA; Department of Pediatrics, University of Iowa, Iowa City, IA, USA

## Abstract

Accurate prognosis after stroke remains challenging, limiting the ability to predict individual recovery. While advanced connectome MRI can provide further prognostically-useful information regarding affected brain networks, its impractical nature has given rise instead to lesion-based estimates of network disruptions. Here, we present a novel pipeline that leverages our recently developed connectome analysis tools to estimate structural disconnection (eSDC) and functional connectivity (eFC) directly from N=96 patient lesion masks for use in predicting motor, language, attention and memory outcomes at multiple timepoints after stroke. Our approach jointly leverages our Network Modification (NeMo) Tool, a database of white matter tractography results that estimates structural disconnectivity (eSDC), and the Krakencoder, a joint autoencoder that estimates functional connectomes (eFC) from the NeMo Tool’s output. Models based on eSDC and eFC predicted almost all outcomes, particularly motor outcomes, significantly better than models that used patients’ fMRI-based observed FC (oFC) and models based on only lesion information and covariates. Estimated connectome measures also i) retained meaningful inter-individual variability when compared to observed functional connectomes, ii) reflected population-level, stroke-related changes in functional connectomes, and iii) resulted in outcome prediction models with feature importances that aligned with expected networks of relevance to the given outcome. Feature importance analyses also identified distinct predictive signatures across both modality and outcome domain, indicating that eSDC and eFC capture non-redundant, biologically meaningful information about post-stroke network disruption. Together, these findings establish our lesion-based estimates of post-stroke structural and functional connectomes as a practical, information-rich alternative to advanced imaging in patients. This approach allows for probing an individual’s affected neural circuitry underlying stroke impairment and recovery, paving the way toward more efficient, accessible, and personalized approaches to post-stroke prognosis and intervention.

## Introduction

Stroke-related brain lesions can result in a range of impairments and affect various functional domains. This complex and highly individual nature of lesion effects and stroke-induced deficits therefore presents a significant challenge in post-stroke rehabilitation. Patients exhibit different brain imaging signatures and recovery patterns, with some but not all showing proportional improvement to their initial impairment and the effectiveness of interventions can vary greatly among individuals.^1–4^ The timing of treatment also significantly influences recovery outcomes, leading to more substantial recoveries in impaired skills.^5^ With stroke the third leading cause of death and disability worldwide and 94 million people living with its effects,^6^ identifying the factors that contribute to individual differences in impairment and recovery, as well as having an accurate prognosis in the acute stage, is crucial for developing tailored treatment and rehabilitation strategies geared toward optimizing recovery.

Advances in brain imaging, such as diffusion magnetic resonance imaging (dMRI) and resting-state functional MRI (rs-fMRI), have facilitated a deeper understanding of how stroke lesions impact large-scale white matter connections quantified via the structural connectome (SC) and their influence on the flow of functional activation represented by the functional connectome (FC), respectively.^7–10^ Our lab, among others, has shown that quantifying properties of the brain’s SC and FC networks and their interrelationship can inform our understanding of brain circuitry underlying behavior in health as well as impairment and recovery in disease, including after stroke.^11–16^ *In silico* simulations of lesions’ impact on structural networks predict widespread disruption of neural dynamics^17,18^ consistent with empirical findings in stroke patients showing that focal damage produces remote FC changes that track behavioral outcomes.^19,20^ It is imperative to understand how lesions disrupt brain networks in order to grasp the global repercussions of stroke damage, resulting impairment, and recovery trajectories.

The development of post-stroke prognoses largely remains dependent on subjective assessments by clinicians, leading to potentially variable predictions and biased allocation of treatment resources.^21–23^ Therefore, there is a critical need for quantitative evidence-based tools to improve prediction precision. Despite numerous prediction tools that have been developed using post-stroke SC or FC and applying mathematical, statistical or machine learning-based models,^24,25,1,7–9,26–28,10^ estimation of stroke outcomes remains difficult due to the substantial inter-individual heterogeneity in lesion topography, network disruption, and post-stroke deficits. Current models are limited in that they have only achieved a moderate level of agreement with observations, lack a focus on preserving inter-individual brain network differences, and lack utilization of multi-modal connectomes (both SC and FC). Compounding this challenge, acquiring advanced dMRI or fMRI imaging to prognose every stroke patient is impractical as long scan times are often poorly tolerated by patients and images are resource- and expertise-intensive to acquire and process. Critically however, most suspected acute stroke patients undergo a short duration, clinical anatomical MRI for diagnostic purposes, that can be in turn used to delineate the stroke lesion, i.e. create a lesion mask. Using information about a lesion’s size and location can enhance prognosis development; particularly useful are techniques that bootstrap lesion masks to estimate their impacts the brain’s networks. Existing lesion-based approaches vary in their capacity to capture network-level disconnection. Lesion-behavior mapping (LBM)^24^ statistically associates lesion location with behavioral deficits but is limited to capturing local anatomical damage, without any network-level information. Lesion-network mapping (LNM)^29,30^ and tractography based structural disconnection tools (BCB Toolkit,^31^ Network Modification (NeMo) Tool^32^) improves on this by mapping lesion location onto a normative functional or structural database to infer the brain networks likely impacted by the lesion. Several previous studies from our group and others have shown that these network disruption estimates are predictive of baseline impairment and future outcomes across stroke and other lesion cohorts^33,34,9,8,7,28,35,36,24,37^. Recently, van den Heuvel et al. challenged the use of LNM as a widely accepted framework to study brain networks underlying neurological disorders, claiming that the networks involved in LNM were largely reflecting nonspecific properties of normative connectomes.^38^ A response to this argument details the lesion-deficit specificity of LNM and claims that a multitude of papers that use LNM do not use the same procedures criticized in van den Heuvel et al., discouraging the invalidation of LNM methodology.^39^ In any case, these lesion-based approaches in question are unimodal by nature and are thus biased toward capturing the effects of either gray matter (LNM) or white matter lesions (structural disconnection tools), and therefore lack a multi-modal estimate of the lesion’s impact on both structural and functional brain networks.^36^ Closing the gap between the clinical practicality of lesion masks and the network-level insight of advanced connectome imaging is essential for accurate brain-behavior mapping, which could help identify relevant networks for modulation with targeted interventions to improve post-stroke outcomes.

Our lab’s recent work in multi-modal connectome mapping has resulted in a tool called the Krakencoder,^40^ which is a joint autoencoder framework with a shared latent space that allows mapping between SC and FC. The Krakencoder’s mapping preserves inter-individual differences in SC and FC in both healthy and lesion cohorts, and provides a low-dimensional, higher signal-to-noise ratio (SNR) representation of multi-modal connectome information. The Krakencoder’s multi-modal connectome representation has also been shown to outperform original connectomes in predicting complex behavior and classifying sex.^40,41^ One of our most recent works used the NeMo Tool to create estimated structural disconnectivity (eSDC) from lesion masks, which was in turn fed to the Krakenocder to create a corresponding estimated FC (eFC). We subsequently showed that the eSDCs and eFCs had moderate to high levels of accuracy in predicting baseline cognition and 4-year cognitive outcomes in a large cohort of people with multiple sclerosis. ^34^

We hypothesized that meaningful multi-modal connectome disruption information can be extracted from clinically-acquired post-stroke lesion masks to accurately predict impairment and outcomes at a level (at least) comparable to models based on observed patient fMRI data in the form of functional connectomes. To test this, we used the previously described NeMo+Krakencoder pipeline to estimate structural and functional connectomes directly from clinically available lesion masks of stroke patients (Fig. 1). To compare our lesion-based estimates of network disruption (eSDCs and eFCs) to observed network disruption, observed functional connectivity (oFC) was extracted from each stroke subject’s acutely collected fMRI. Statistical models were then used to predict motor, language, attention, and memory outcomes in stroke patients collected at the acute, 3-month, and 1-year timepoints using estimated or observed connectome information. If our models based on estimated network disruptions performs equally well compared to models based on observed networks, this would support the use of clinical MRI in lieu of advanced MRI for accurate post-stroke prognosis development and identification of potentially targetable networks to enhance post-stroke outcomes.

**Figure 1:**
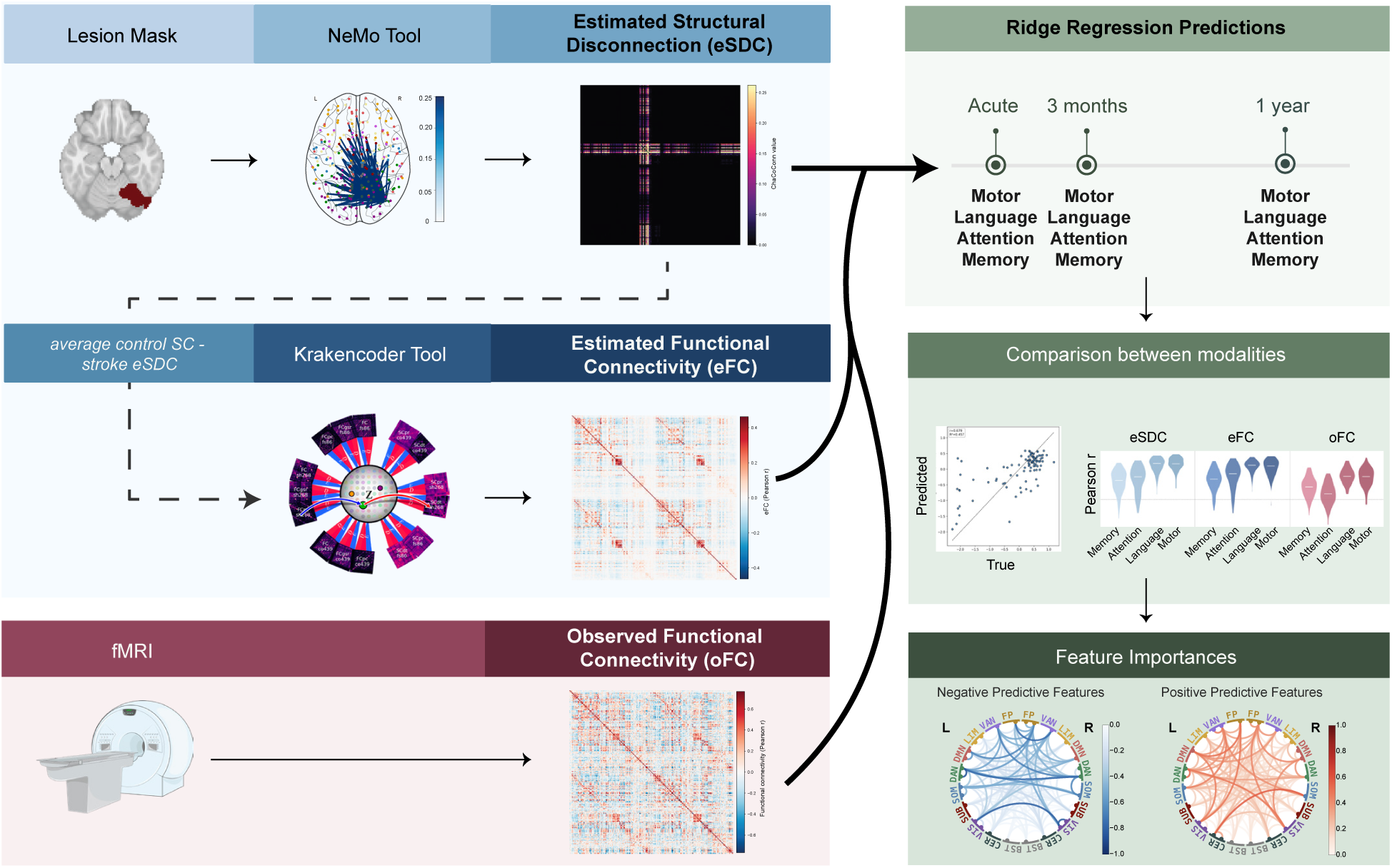
Workflow. Lesion masks from clinical MRI in stroke patients are used to estimate structural and functional connectomes. The NeMo Tool quantifies streamlines intersecting the lesion from a normative tractography database to generate estimated structural disconnection (eSDC). The Krakencoder, a joint autoencoder framework, maps between structural and functional connectivity and is used to estimate functional connectivity (eFC) from the NeMo Tool’s eSDC. Estimated connectomes are then used to predict post-stroke outcomes and compared against models using observed functional connectivity (oFC) derived from patient resting-state fMRI and covariate only models. Model performances are statistically compared and model feature importance extracted.

## Results

We used a cohort of 132 adults admitted to the Washington University School of Medicine who had ischemic or hemorrhagic stroke anywhere in the brain that resulted in clinical evidence of post-stroke deficits.^25^ Scans include both anatomical (T1, voxel size = 1.0 x 1.0 x 1.0 mm) and rs-fMRI (duration up to 30 mins across several 4m16s runs, TR = 2,000 ms, voxel size = 4 x 4 mm) taken at the acute (*∼*2 week) post-stroke timepoint. All patients underwent a battery of tests that were collected at the acute, 3-month, and 1-year timepoints post-stroke and included verbal memory, visual memory, left/right attention, language, left/right upper motor and left/right lower motor function measures (See Methods for score descriptions). MRI scans were also collected at the 3-month and 1-year time points, but they were not utilized here as the focus was on using baseline (acute) imaging to predict future impairment and outcomes. It has been previously shown that post-stroke structural disconnection estimated using the NeMo Tool is significantly and moderately associated with structural connectivity changes derived from patient-based dMRI tractography.^42^ Thus, in this work, we focus on the novel extension of comparing our lesion-based estimated FC to observed FC, and comparing the post-stroke outcome prediction accuracy of our multi-modal estimated and observed connectome data. After MRI quality control and accounting for missing behavioral or covariate data, the final stroke patient cohort was acute N=96 (aged 19-77; 43.8% female), 3-months N=78 (aged 19-77; 44.9% female), and 1-year N=70 (aged 19-77; 44.3% female). Subjects vary by outcome and timepoint because not all participants completed every outcome assessment at each timepoint visit. Supplementary Table 1 reports outcome descriptive statistics and sample sizes for each time point. Key clinical and demographic information for the final cohort is summarized in Table 1. For comparison, a cohort of N=30 non-stroke controls was enrolled and underwent two MRI scans 3 months apart, including resting-state fMRI, anatomical scans, diffusion MRI, and the same neuropsychological battery (N=22 after MRI quality control; aged 21-83, 47.6% female).^25^

**Table 1:** Demographic and clinical characteristics of the stroke cohort (N = 96). Values are mean *±* SD (range) for continuous variables and N (%) for categorical variables. NIHSS: National Institutes of Health Stroke Scale. NIHSS data were available for 86 participants.

| **Characteristic** | **Value** |
|---|---|
| Age, years | 52.1 ± 9.7 (19–77) |
| Sex |  |
| Female | 42 (43.8%) |
| Male | 54 (56.2%) |
| Handedness |  |
| Left | 9 (9.4%) |
| Right | 87 (90.6%) |
| Race |  |
| Black or African American | 66 (68.8%) |
| White | 30 (31.2%) |
| Ethnicity |  |
| Not Hispanic or Latino | 96 (100.0%) |
| Education, years | 13.0 ± 2.4 (9–20) |
| Lesion type |  |
| Ischemic | 77 (80.2%) |
| Hemorrhagic | 13 (13.5%) |
| Other | 6 (6.2%) |
| Lesion side |  |
| Left | 50 (52.1%) |
| Right | 46 (47.9%) |
| Lesion site |  |
| Subcortical | 16 (16.7%) |
| Cortical | 33 (34.4%) |
| Cortico-subcortical | 22 (22.9%) |
| White matter only | 5 (5.2%) |
| Brainstem | 10 (10.4%) |
| Cerebellar | 7 (7.3%) |
| Other | 3 (3.1%) |
| Lesion volume, cm<sup>3</sup> | 33.25 ± 48.07 (0.1–277.0) |
| NIHSS | 7.0 ± 6.1 (0–25) |
| Days from stroke to acute scan | 13.6 ± 4.5 (6–28) |

### Observed and lesion-estimated post-stroke connectome disruptions are correlated

The stroke cohort lesion locations are visualized via a heatmap in Figure 2a. Individual patients’ MNI space lesion masks were entered into the NeMo tool to estimate structural disconnectivity matrices.^32^ The NeMo Tool uses a normative tractography database from 420 healthy controls (unrelated individuals from the Human Connectome Project-Young Adult (HCP-YA) dataset; *∼*50% female, aged 22-36) and quantifies for each SC edge (pair of regions in an atlas), the percent of its white matter streamlines that pass through the lesion. The result is an eSDC matrix per stroke subject, where a value of 1 indicates a complete disconnection, and a value of 0 indicates no disconnection. The group average eSDC across the stroke cohort is visualized in Fig. 2b. Edges are arranged by network, where the networks include the Yeo-7^43^ definitions (frontoparietal (FP); ventral attention (VAN); limbic (LIM); default-mode (DMN); dorsal attention (DAN); somatomotor (SOM); visual (VIS)), along with subcortex (SUB), cerebellum (CER), and brainstem (BST). The NeMo Tool also outputs the corresponding estimated SC (eSC), which is calculated by counting the number of streamlines between pairs of regions after streamlines passing through the lesion are removed; these are retained for use in the Krakencoder’s estimates of FC (see below).

**Figure 2:**
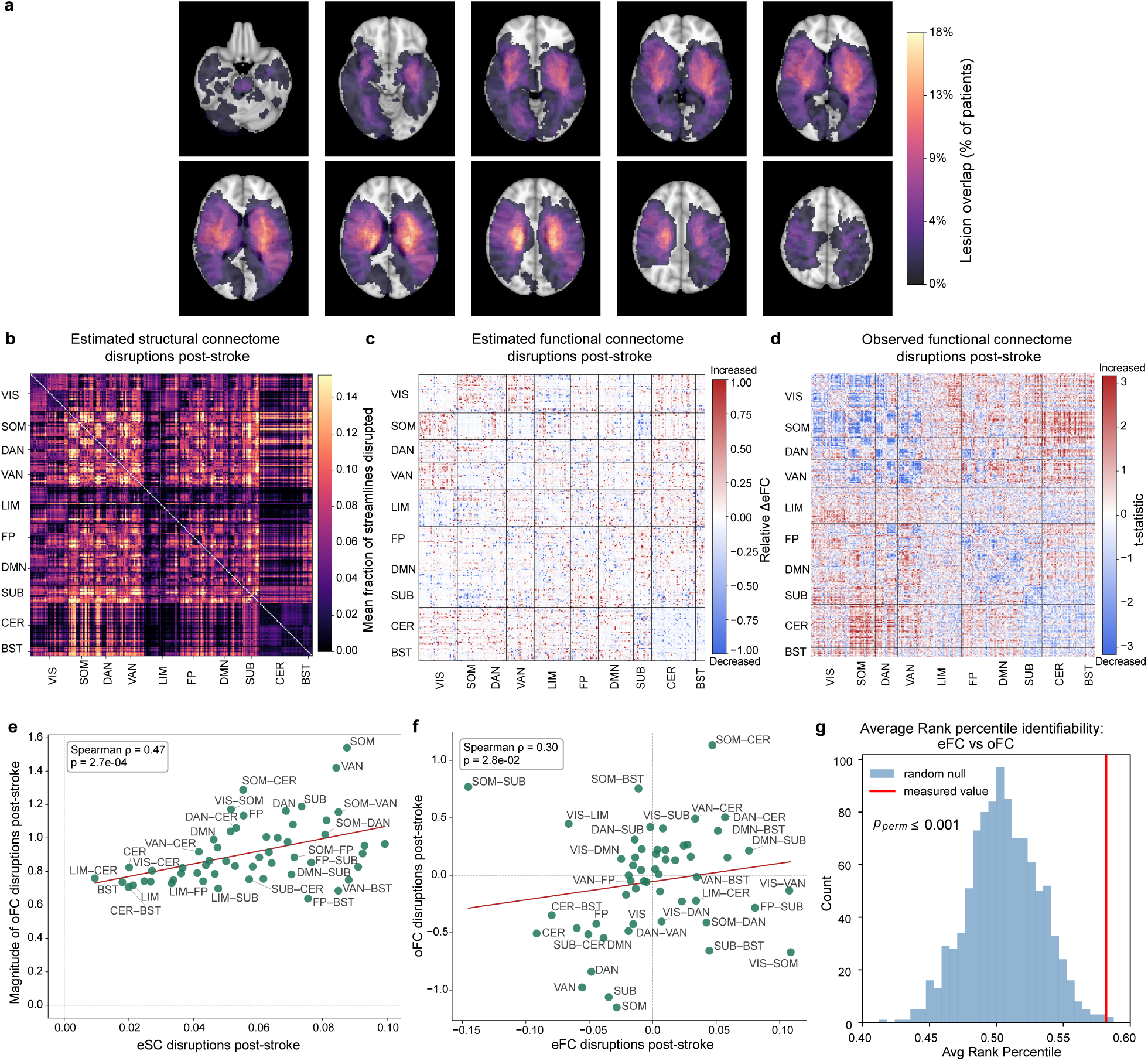
Structural and functional connectivity disruptions following stroke. **a)** Stroke lesion location heatmap (N=96) in MNI space. **b)** Estimated structural disconnectivity (eSDC) matrices, averaged across 96 stroke subjects. Regions are ordered by network assignment (VIS - visual, SOM - somatomotor, DAN - dorsal attention, VAN - ventral attention, LIM - limbic, FP - frontoparietal, DMN - default mode network, SUB - subcortical, CER - cerebellum, BST - brainstem). **c)** Estimated functional connecomte (eFC) disruptions post-stroke. Values are relative % difference between the stroke cohort’s mean eFC and the mean observed FC from NeMo Tool controls (N=420). **d)** Observed functional connectome (FC) disruptions post-stroke. T-statistic comparison of observed stroke FC (N=96) vs observed FC of the stroke study’s controls (N=22). **e)** estimated structural disconnection (as in panel b) vs observed FC disruptions (*|*oFC *t*-statistic*|*) (as in panel d), presented as network-pair averages (labels indicate the network pair, and within network values have one label). **f)** estimated FC changes (as in panel c) and observed FC changes (raw *t*-statistic, as in panel d), presented as network-pair averages. **g)** Average rank percentile for identifiability of the observed and estimated FC for the stroke cohort (red line) vs permutation-based null (blue histogram).

Next, the stroke individual’s eSC is fed into the Krakencoder tool, a joint autoencoder with a shared latent space that maps between structure and function, to output eFCs for every individual with stroke. To visualize the average estimated FC disruption caused by stroke, we calculated the per-edge relative difference between the mean eFC from the stroke cohort and mean observed FC from the 420 HCP-YA controls used in the NeMo Tool; see Fig. 2c. Relative differences were calculated per edge by subtracting the stroke cohort’s mean eFC from the 420 HCP-YA controls’ mean oFC and dividing this by the mean of the two values. We did not apply a t-test here as the stroke cohort’s eFCs are derived from the HCP-YA data and are thus not independent; instead we wished to quantify how the stroke eFCs are changing from the reference set of HCP-YA oFCs. Finally, we calculated the per-edge t-statistic of the oFCs from the stroke cohort compared to the healthy age-matched controls (N = 22) that had identical fMRI acquisitions; see Fig. 2d. This allows quantification and visualization of observed FC disruptions occurring in our cohort post-stroke, contrasted against a comparable set of normative FCs.

We tested our hypothesis that estimated structural and functional disruptions due to stroke would be associated with observed functional disruptions, at the level of the entire stroke cohort. To do this, we correlated the network-pair averages of i) the stroke cohort’s mean eSDC (Fig. 2b) and ii) the absolute value of the oFCs’ mean t-statistics (stroke vs healthy controls; Fig. 2d); see Fig. 2e. We used the absolute mean t-statistics to quantify the magnitude of oFC disruption, irrespective of whether FC was increased or decreased relative to controls. Stroke-related eSDCs are positively correlated with the disruption in oFCs (Spearman *ρ* = 0.47, *p* = 2.7*e^−^*^4^), indicating that network pairs with greater estimated structural disconnection also exhibited larger deviations (regardless of direction) in observed FC from healthy controls. Similarly, we correlated the network-pair averages of i) the stroke cohort’s mean eFC relative difference (eFC vs HCP-YA oFC; Fig. 2c) and ii) the mean oFCs’ t-statistics (stroke vs healthy controls; Fig. 2d); see Fig. 2f. Here we took the signed mean t-statistic, as the eFC should reflect the directionality of the stroke related changes in oFC. These were also positively correlated (Spearman *ρ* = 0.30, p=0.028), indicating that, after stroke, network pairs with increases/decreases in lesion-based estimates of FC also tended to have increases/decreases in fMRI-based observed FCs relative to healthy controls. We also examined these associations at the individual level. The mean absolute oFC z-score across edges within each network pair was significantly correlated with the corresponding mean eSDC (Spearman *ρ* = 0.16, p = 0.0028; Supplementary Fig. 1a). In contrast, the mean oFC z-score across edges within each network pair was not significantly correlated with the corresponding mean relative difference in eFC (Supplementary Fig. 1b).

We next conducted an identifiability analysis to directly compare eFCs and oFCs for each stroke patient, asking whether a person’s lesion-based eFC is more similar to their own fMRI-based oFC than other stroke subjects’ fMRI-based oFCs. This was done by calculating and then ranking the Pearson correlation between an individual’s eFC and the entire group of stroke subjects’ oFCs. The rank of the correctly matching individuals’ oFC was computed and summarized as the average rank percentile. A value of 100% indicates a person’s own oFC is always ranked first (perfect identifiability) when compared to the entire group of oFCs. To gather a distribution of chance values of the average rank percentile, which should be *∼*50%, the subjects’ oFCs were permuted and the average rank percentile was calculated (see Fig. 2g). The average percentile for our data was 58% (*p*_perm_ *<* 0.001) revealing that the Krakencoder’s eFCs are slightly but significantly more similar to each patient’s own oFC than to other patients’ oFCs. This result was surprising; it suggests that our estimated FC, derived solely from lesion masks and without additional subject-specific information, captures some of the individual-level variation in post-stroke individuals’ fMRI-based, observed FCs.

Together, these findings reveal that our estimated measures (eSDCs, eFCs) to a certain extent reflect observed functional post-stroke connectome disruption. Additionally, although eFCs are derived solely from lesion masks via the NeMo tool and the Krakencoder, with no resting state fMRI involved, they still capture enough subject-specific information to identify a subject’s own fMRI-based oFC above chance levels. This finding is significant as it validates our pipeline as a useful tool to bootstrap clinically-acquired lesion masks to estimate how functional and structural connectomes are disrupted by lesions.

### Lesion-estimated connectivities outperform observed functional connectivity in predicting post-stroke outcomes

We used ridge regression with nested cross validation to predict visual memory, verbal memory, left/right visual attention, language, left/right upper motor, and left/right lower motor composite scores at the acute, 3-month, and 1-year timepoints post-stroke. Prediction accuracy was compared among the eSDC, eFC, and oFC models, as well as a covariate-only baseline model (called COV) that included lesion volume, lesion side, age, sex, and days from stroke to scan; see Fig 3. All models based on connectome disruption metrics (eSDC, eFC, oFC) include the variables in the COV model. Overall, the majority of acute outcome model accuracies (8 out of 9) were significantly higher for estimated connectome models (eSDC and/or eFC) compared to oFC and/or covariate models. This pattern persisted for 3 month outcome prediction, with 7 out of 9 outcomes having the highest accuracy for estimated connectome models and performing comparably to or significantly better than oFC and/or COV models. At the 1 year time point, estimated connectome models outperformed the covariates and/or oFC models in only the motor outcomes (4 out of 9). Motor outcomes were the most predictable in general, and both eSDC and eFC predicted significantly better than oFC and covariate models across all timepoints for left/right upper motor and left lower motor models (combined median r range 0.45–0.79). Only 8/27 models (visual memory at all timepoints, 3-month verbal memory, 1-year left/right attention, verbal memory and language outcomes) were significantly less predictable with estimated connectomes compared to oFC and/or covariate models. See Supplementary Fig. 2 for full Pearson r prediction distributions for all outcomes.

**Figure 3:**
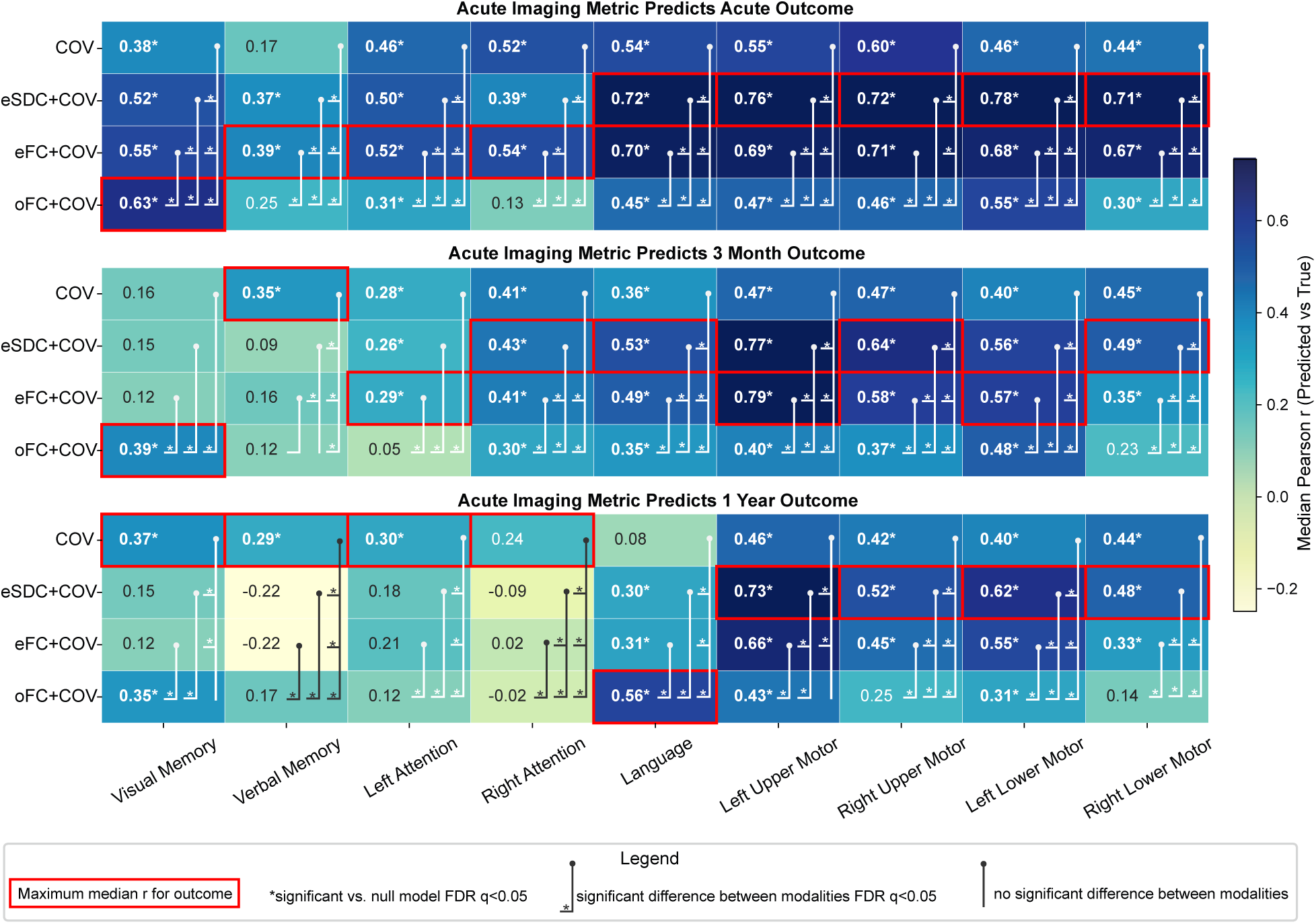
Prediction of post-stroke outcomes. Median Pearson *r* between predicted and observed outcome scores for the test set. Models based on the four different data modalities (eSDC, eFC, oFC, and a covariate only model - COV) are shown in each row, with a given post-stroke outcome domain in each column, and post-stroke timepoint by panel. All imaging models contain the covariates in the COV model. Asterisks after the median Pearson *r* value indicate models whose performance significantly exceeded a permutation-derived null model (permutation test; FDR *q <* 0.05). Brackets with left tickmarks indicate significant differences in that model’s performance compared to the model where the bracket begins, denoted with an asterisk (permutation-based Wilcoxon rank-sum test on fold-wise Pearson *r* values; FDR *q <* 0.05). Red boxes highlight outcomes with the highest median prediction performance across modalities.

To examine whether sex influenced the prediction models, we trained models separately on male and female participants and tested each on the same held-out male and female participants, with training and test set sizes matched across sexes (Supplementary Table 2). Interestingly, models trained on female data tended to be better in predicting male outcomes than models trained on male data (Supplementary Fig. 3). Due to the small number of participants per sex, this analysis was exploratory.

### Brain-behavior maps relating connectomes to post-stroke outcomes vary by modality

We investigated which brain network connections are driving the predictions for different modalities and outcomes. We hypothesized that although eSDCs and eFCs are both derived from lesion masks and had similar accuracies for most outcomes, the feature importances contributing to these predictions would vary. We used the Haufe transformation^44^ on the models’ beta coefficients in order to better capture a biologically interpretable feature importance map (see Methods). For each modality and outcome model, region-pair coefficients’ significance was assessed within each of the 500 outer cross-validation folds (5-fold cross-validation repeated 100 times). For each connectivity model coefficient per fold, a one-sided permutation p-value was computed as *p* = (*n_exceed_* + 1)*/*(*n_perm_* + 1), where *n_exceed_* is the number of permutations in which the absolute null-permuted beta weight was greater than or equal to the absolute observed beta weight. A connection was considered significant in a fold if uncorrected p *<* 0.05, and connections with beta weights that were significant in at least 80% of folds were retained. This was not corrected for multiple comparisons as it was a stability criterion. For each retained connection’s beta weight, the Haufe-transformed feature importance was averaged across all folds to obtain a single feature importance value. Within each pair of hemisphere-specific networks, positive and negative Haufe values of retained edges were then averaged separately. Figure 4 shows examples of the positive and negative feature importances driving the predictions for acute left lower motor (Fig. 4a), language (Fig. 4b), and visual memory (Fig. 4c). A negative predictive feature (blue chord) indicates that an increase in connectivity (or an increase in disconnection for eSDC) between the two networks predicts a worse score, and a positive predictive feature (red chord) indicates that an increase in connectivity (or an increase in disconnection for eSDC) predicts a better score, with color indicating magnitude. Significant feature importances for all other outcomes at the acute timepoint can be found in Supplementary Fig. 4.

**Figure 4:**
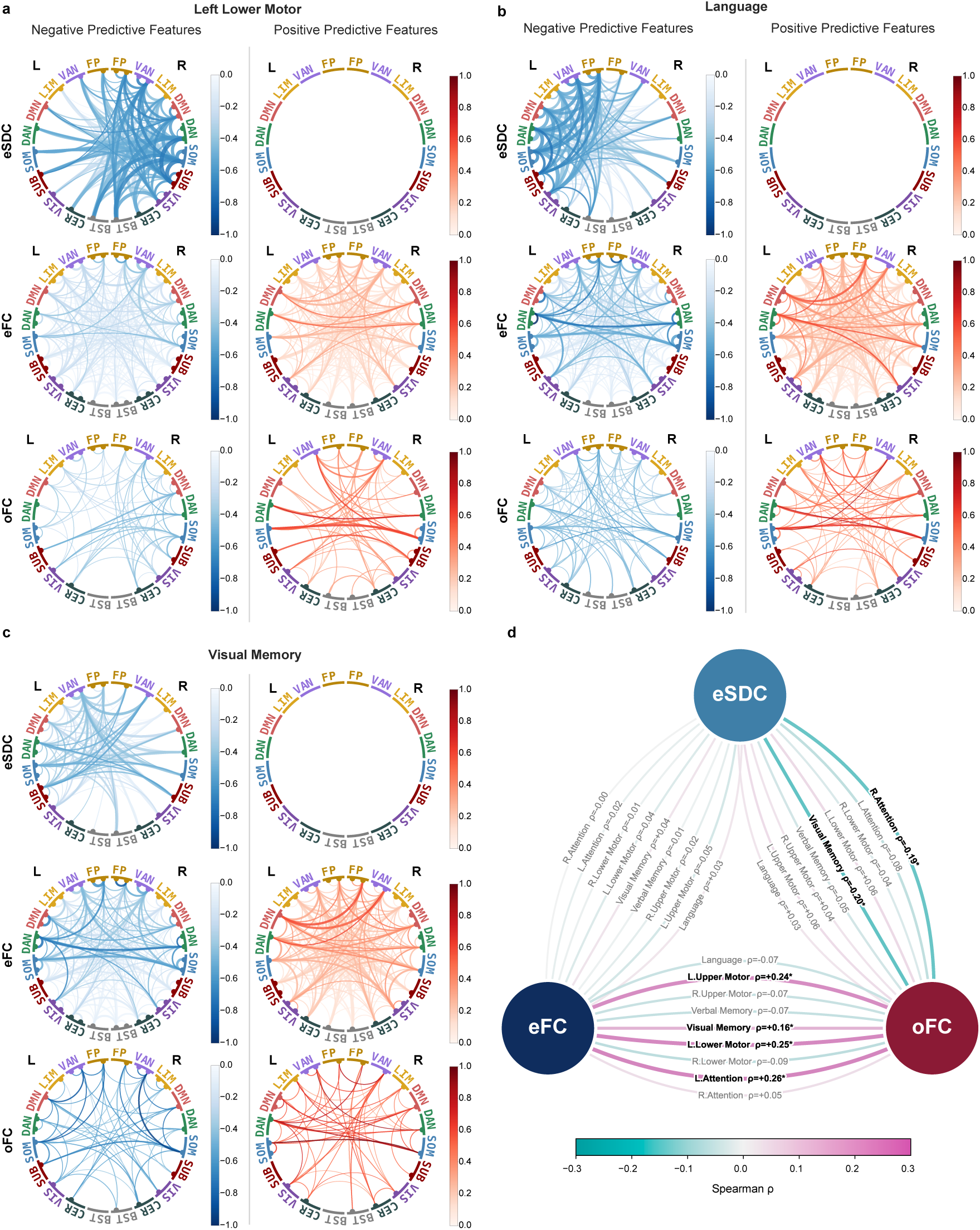
Network-pair feature importances for models predicting acute poststroke outcomes. Network-level feature importances driving predictions for left lower motor (**a**), language (**b**), and visual memory (**c**) outcomes at the acute timepoint, shown separately for eSDC, eFC, and oFC models (top, middle and bottom, respectively). Chord plots show network-pair averages of significant positive predictive features (hot colorbar) where larger (dis)connectome values indicate better behavioral scores and significant negative predictive features (cool colorbar) where larger (dis)connectome values indicate worse behavioral scores. Chord width is proportional to the square root of the fraction of edges in each network pair that met the stability criterion with that sig^1^n^5^, on a common scale across all panels. L and R denote left and right hemispheric networks. **d** Between-modality Spearman correlation of all acute outcome models’ network-level feature importances; *Uncorrected permutation p<0.05.

Significant negative eSDC features (more disconnection = worse outcomes) were strongly lateralized to the right hemisphere for left lower motor predictions and to the left hemisphere for language predictions, as expected. Negative features for eSDC were also lateralized to-wards the left hemisphere for visual memory predictions, with noticeable gaps from the left BST and left CER compared to left lower motor and language. There were no significant positive eSDC features were across outcomes, also as expected. Contrastingly, significant eFC and oFC feature importances were more distributed throughout both positive and negative connections across outcomes.

As an exploratory analysis to test our hypothesis that each modality has unique feature importances contributing to predictions, we correlated average network-level Haufe-transformed feature importances between modality pairs within outcomes at the acute time-point (Fig. 4d). eFC and oFC patterns were positively correlated for left upper motor (Spearman *ρ* = 0.24), left lower motor (*ρ* = 0.25), left attention (*ρ* = 0.26) and visual memory (*ρ* = 0.16), with all uncorrected p *<* 0.05 and corrected p *>* 0.05. eSDC and oFC patterns were negatively correlated for right attention (*ρ* = -0.19) and visual memory (*ρ* = -0.20; p ¡ 0.05, uncorrected). eFC features did not significantly correlate with eSDC patterns for any outcome tested, indicating that Krakencoder-derived eFC relies on different network connections than its structural input to generate predictions.

To test whether network-level feature importances remained stable when predicting outcomes at later timepoints, we correlated feature-importance patterns within each modality across timepoints, but only for models that significantly exceeded their permutation null. Network-level feature importances were highly stable across timepoints for eSDC and eFC (median *ρ* = 0.991 and 0.983, respectively), and somewhat less so for oFC (median *ρ* = 0.801; range 0.389–0.978; Supplementary Table 3). These results indicate that the Haufe patterns reflect outcome-specific information that remained consistent across 3-month and 1-year predictions.

## Discussion

In this work, we use a novel pipeline to obtain estimated structural and functional connectomes (eSDCs and eFCs) derived from stroke lesion masks to predict motor, language, attention, and memory outcomes at three timepoints across recovery. Specifically, we used the NeMo tool to estimate the structural connectivity disruptions (eSDC) of stroke lesions without requiring tractography in stroke patients. The Krakencoder was then used to estimate FC directly from the eSDCs, extending our pipeline’s reach from structural disconnection alone to a full multi-modal connectome estimate - all without requiring any advanced imaging beyond the original, clinically acquired lesion mask. Remarkably, we find the NeMo and Krakencoder pipeline for estimating FC preserves individual-level features present in the observed FC derived from patient fMRI data. Further, we find that outcome prediction models based on estimated connectomes largely outperform models based on observed FCs and covariate-only model predictions. Finally, we find outcome prediction models based on estimated structural and functional connectomes, despite being derived from the same lesion masks, capture distinct feature importance signatures. Altogether, these results validate a framework for efficient extraction of multi-modal connectome disruption information using only lesion masks, which in turn can be used to predict recovery trajectories and isolate brain networks relevant to a given outcome - all without the use of advanced MRI.

Estimating connectome disruptions from lesion masks provides a clinically feasible way of bootstrapping readily available information to create more accurate patient prognoses. While advanced MRI is difficult to acquire after stroke, costing time and money, it has been unclear whether lesion-based estimates of connectome disruption track an individual’s actual network damage or only the average effect of lesion location on a normative brain. Here, network-level disruption estimated by eSDC and eFC was significantly correlated with functional disruption measured directly in each patient’s fMRI-based observed FC (eSDC: Spearman *ρ*=0.47, p=2.7×10^-4^; eFC: *ρ*=0.30, p=2.8×10^-^²), indicating that our lesion-based connectome estimates capture stroke-related connectome disruption rather than an arbitrary approximation. Remarkably, our estimates of post-stroke FC also preserved identifiability information: each patient’s estimated FC was more similar to their own observed FC than to any other patient’s observed FC. To our knowledge, this is the first demonstration that estimated functional connectivity is identifiable at the level of the individual patient after stroke. This suggests that lesion-derived estimates via the Krakencoder recover individual-level signal rather than a shared group-average pattern, and motivates their continued development for clinical use.

While our lesion-based estimates of connectivity (eSDC and eFC) were significantly related to observed FC, our estimated models generally significantly outperformed models based on observed FC and models based only on covariates when predicting post-stroke outcomes. Models based on estimated connectomes especially performed well across all time-points when predicting motor outcomes. Previous work with this same stroke cohort used LNM to derive functional and structural disconnection maps (FDC and SDC). It found that FDC maps predicted worse than fMRI-derived FC across outcomes, and that SDC maps predicted impairment about as well as lesion information alone.^28^ We find a different result with our NeMo-derived eSDC and NeMo+Krakencoder-dervied eFC models by showing their accuracy actually exceeds that of models based on fMRI-derived oFC and lesion information+covariates. LNM was developed primarily to identify the shared networks to which lesions in different locations map when they produce the same symptom, rather than to predict individual scores.^33,30,39^ We therefore view our results as complementary to the insights LNM provides, and as an exciting step toward more accurate prediction of post-stroke outcomes using lesion masks and uncovering the individualized networks contributing to those predictions.

Using our models mapping connectomes to outcomes, we examined the specific brain network connections driving these predictions by deriving network-level Haufe-transformed feature importances. Feature-importance patterns were largely distinct across connectome modalities. Correlations were significant between eFC and oFC for left upper and lower motor, left attention, and visual memory (*ρ* = 0.16-0.26, p*<* 0.05; none survived correction for multiple comparisons), suggesting partial overlap in the networks contributing to predictions of these outcomes; interestingly, three of the four were left-sided outcomes, which depend predominantly on right hemisphere connections. eSDC and oFC patterns were negatively correlated for right attention (*ρ* = *−*0.19) and visual memory (*ρ* = *−*0.20; uncorrected p *<* 0.05), such that network pairs in which greater structural disconnection was associated with worse outcomes tended to be those in which stronger functional connectivity was associated with better outcomes. Notably, eFC feature importances did not correlate with eSDC’s at all, despite eFC being derived directly from eSDC through the Krakencoder tool. This suggests that eSDC and eFC are not redundant estimates of the same signal, but instead capture unique aspects of post-stroke network disruption, each distinct to different outcomes.

Inspecting the models’ feature importances allows for a sanity check and validation of our novel pipeline. Negative eSDC features were concentrated in the left hemisphere for right upper and lower motor, right attention, language, and visual and verbal memory predictions, and in the right hemisphere for left upper and lower motor and left attention predictions. In these models, greater structural disconnection involving the affected hemisphere’s networks, within it and with the rest of the brain, was associated with worse predicted scores in a way that aligns with known lateralization of human brain function. For the motor and attention outcomes, this pattern follows the contralateral organization of sensorimotor and attentional control and is further consistent with prior work linking interhemispheric white matter asymmetry to reduced motor function in stroke patients.^45^ For language it follows the left-lateralized perisylvian network and prior evidence linking structural disconnection within dorsal and ventral language pathways to aphasia severity and naming deficits.^46,47^ There were no significant eSDC positive features, with the exception of verbal memory that had one surviving feature low in magnitude, consistent with the expectation that an increase in structural disconnection has little plausible mechanism for improving outcomes. In contrast, significant eFC and oFC features were spread across both hemispheres and networks and included both positive and negative weights, consistent with prior evidence that stroke disrupts widespread functional networks in both hemispheres^48^ and can result in either functional disconnection or upregulation.^49^ Observed FC models generally retained fewer significant features than the estimated FC models. One possible explanation is that the noise in fMRI-based connectivity limits which individual features meet our stability criterion, whereas eFC is more stable and less affected by this noise.

Limitations of this approach include the potential for weak generalizability, given our relatively small dataset of 96 stroke patients all drawn from a single study. Recent work has debated the sample sizes required for reliable brain-behavior mapping; however, a recent analysis using Adolescent Brain Cognitive Development data concluded that sample sizes below 100 are adequately powered for mapping MRI metrics to behavior.^50^ While we use observed, fMRI-based FC as a benchmark throughout this study, FCs are known to be noisy, especially for a single-timepoint measurement and short scan time in a clinical population and thus may not be a good ground truth. Our finding that eSDC and eFC outperform observed FC should therefore be interpreted as evidence of their practical utility and robustness, not as evidence that they are superior in any absolute sense. Sex-specific analyses were limited by small per-sex samples (n = 27-40) and should be considered exploratory; larger cohorts will be needed to determine whether brain–behaviour relationships after stroke differ by sex. Finally, our lesion-based estimates are (by design) biased toward representing the lesion effects in a static way, and thus (unlike possible fMRI-based FC) will not capture any functional reorganization that may happen in the recovery phase after stroke^9^.

We demonstrate that our novel pipeline for estimating functional and structural network disruptions from lesion masks can provide a time-efficient and clinically feasible yet behaviorally relevant alternative to performing advanced connectome MRI in patient populations. This project addresses a major gap in the current state of the field by providing personalized insights into the neural mechanisms underlying post-stroke impairments and recovery. The ability to more accurately and efficiently predict outcomes could guide rehabilitation efforts and help patients and their providers more effectively plan for the level of services that may be required. For example, patients identified as being at risk for long-term motor impairment may be identified and more aggressive treatment plans created for those patients. The findings from this project will lay the groundwork for identifying individual variability in connectome disruption that is related to post-stroke impairment outcomes, which is a crucial first step in identifying targets for therapeutic intervention. Future work could focus on analyzing treatment effects in post-stroke cohorts, e.g. transcranial magnetic stimulation studies,^51^ using insights from the models built here. The collection of larger and more diverse datasets, especially those that integrate information related to sex-differences, genetic, and clinical information, could enhance the Krakencoder model’s predictive capabilities and provide a more comprehensive understanding of the connectome’s role in stroke recovery. Validating our models’ predictions in independent cohorts and exploring their utility in clinical settings could help translate these findings into personalized treatment strategies for stroke patients.

## Methods

### Participants

The data used in this project is from a cohort of 132 stroke patients that were admitted to the Washington University School of Medicine (WUSM). Patients underwent both anatomical and resting state functional MRI and a battery of neuropsychological tests. Participants were excluded from our analyses if required behavioral measures, covariates, lesion masks, or quality-controlled imaging data were unavailable. A final cohort of N=96 (aged 19-77; 43.8% female) stroke subjects remained after these quality controls. A healthy control group (N=30; N=22 after quality control, aged 21-83, 47.6% female) was matched with the study sample for age and sex; they also underwent the same MRI and battery testing at two visits 3 months apart.^25^

### Outcome measures

Behavioral outcomes were assessed at the acute, 3 months and 1 year timepoints, and grouped into domains. **Visual memory:** Brief Visuospatial Memory Test^52^ (BVMT) immediate recall, BVMT delayed recall, Spatial span forward, Spatial span backward.^53^ **Verbal memory:** Hopkins Verbal Learning Test^54^ (HVLT) immediate recall, HVLT delayed recall, HVLT discrimination index. **Left/right attention:** Visual field accuracy valid/invalid, Reaction time valid/invalid,^55^ Mesulam omissions^56^, Behavioral Inattention Test (BIT) omissions.^57^ **Language:** Word reading, Commands, Boston Naming, Complex ideational material, Reading, Reading comprehension^58^ **Left/right upper-limb motor function:** Shoulder flexion, Wrist extension,^59^ Grip strength,^60^ 9-hole peg test,^61^ Action Research Arm Test.^62^ **Left/right lower-limb motor function:** Ankle flexion, Knee extension, Hip flexion, Ankle motor strength, Knee motor strength, Hip motor strength.^59^

For each domain, the outcome was a composite: each measure was z-scored using the mean and standard deviation of the training set in each cross-validation fold, and the z-scores were averaged. Measures for which lower values indicate better performance (reaction times and omissions) were sign-reversed before averaging, so that higher composite scores indicate better performance. Participants were included for a domain at a given timepoint if all of its measures were available.

### fMRI acquisition, preprocessing, and postprocessing

Patients underwent anatomical (T1/T2, 1mm voxels) and rs-fMRI scans at the acute ( 2 week) timepoint post-stroke (mean = 13.6 days, SD = 4.5 days) at WUSM. All scans were acquired on a Siemens 3T Tim-Trio with a 12-channel head coil and scans were collected using a gradient echo EPI sequence (TR = 2,000 msec, TE = 27 msec, 32 contiguous 4-mm slices, 4×4 mm in-plane resolution) while patients stared at a central cross in a low-light setting. This acquisition resulted in six to eight runs with 128 volumes, or about 30 minutes total, for each patient. Healthy age-matched controls underwent the same scanning protocol. The full MRI protocol is described in Corbetta et al.^25^ T1s were pre-processed using FreeSurfer (v7.3.2) to obtain tissue types and regional parcellations. T1 scans for 6 of the 96 patients had motion artifact or lesion size significant enough to disrupt anatomical processing and spatial normalization to MNI, and these were replaced with synthetic T1 reconstructions using FreeSurfer SynthSR^63^. Functional MRI data from WashU used fMRIPrep (v23.2.1)^64^ for initial preprocessing of BOLD data, including generation of a reference volume and skull-stripped version, estimation of head-motion parameters using mcflirt,^65^ and volume-based spatial normalization to MNI152NLin6Asym. Because no acquired field map was available, a deformation field to correct for susceptibility distortions was estimated based on fMRIPrep’s fieldmap-less approach. The deformation field is that resulting from co-registering the EPI reference to the same-subject T1w-reference with its intensity inverted.^66,67^ Registration was performed with antsRegistration (ANTs 2.5.0),^68^ and the process regularized by constraining deformation to be nonzero only along the phase-encoding direction, and modulated with an average fieldmap template.^69^ The BOLD reference was co-registered to the T1w reference using bbregister,^70^ and confounding time-series were calculated based on the preprocessed BOLD. Post-processing included automatic removal of motion artifacts using independent component analysis (ICA-AROMA)^71^ that was performed on the preprocessed BOLD on MNI152NLin6Asym space time-series after removal of non-steady state volumes and spatial smoothing with a nonlinear filter that preserves underlying structure (6.0 mm FWHM).^72^ This was used only to estimate artifacts to remove; the denoised data was not spatially smoothed. Outlier timepoints were identified as those with framewise displacement^73^ above 0.9mm or global signal above 5*σ*. Global signal regression (removing the global gray matter signal and its temporal derivative) and high-pass filtering *>* 0.008 Hz were applied, with outliers excluded. Automated quality control for all runs included framewise displacement and DVARS thresholds, with individual runs excluded if more than 48 of the 128 volumes were motion or global signal outliers. The first four valid runs were used for each subject, and subjects with fewer than four valid runs were excluded from the analysis.

### Observed Functional Connectivity (oFC)

Observed functional connectivity was computed from the denoised resting-state BOLD time series. Regional time series were extracted by averaging voxel time series in each region of the MNI-space Shen268 volumetric parcellation atlas.^74^ The time series for each run was z-normalized and the normalized time series were concatenated. Region-wise functional connectivity matrices were constructed by computing Pearson correlation coefficients of region averaged outlier-free time series. oFC matrices were generated for each stroke participant at the acute timepoint and healthy control for visit one and/or two if applicable.

### Estimated Structural Disconnection (eSDC)

T1 MRIs were registered to the Montreal Neurological Institute brain using FSL’s FNIRT.^75^ Individual structural MRI images (T1, T2, and FLAIR) were manually segmented with lesions and reviewed by board-certified neurologists.^10^ Lesion masks were entered into the NeMo Tool^32^ to calculate eSDCs for each patient. First, the NeMo Tool identifies and removes streamlines in a reference MNI-space tractography database that pass through the lesion mask. This reference database was generated using deterministic tractography (MRtrix3 SD STREAM^76,77^) from 420 unrelated healthy controls (206 female, 214 male, 28.7 ± 3.7 years) from the Human Connectome Project Young-Adult (HCP-YA) dataset. For each pair of regions in the Shen268 parcellation, eSDC was defined as the fraction of total structural connectivity disrupted by the lesion, calculated as the sum of SIFT2 weights^78^ of lesionintersecting streamlines divided by the sum of SIFT2 weights of all streamlines connecting that region pair, yielding 268 x 268 matrices for each subject with a diagonal of zero. Regional eSDC was calculated as the sum of the columns in the eSDC matrix.

### Estimated Functional Connectivity (eFC)

In addition to outputting eSDCs, the NeMo tool generated estimated structural connectomes for each of the stroke patients, calculated by counting the number of streamlines between pairs of regions after streamlines passing through the lesion are removed. This estimated SC was used as input to the Krakencoder^40^, a joint autoencoder with a shared latent space that simultaneously and bidirectionally maps between various flavors of SC and FC across different atlases and processing choices. Here, we use the Krakencoder to map from NeMo-estimated SC to an estimated FC. To ensure comparability with the observed oFCs used in this study, eFC was generated using the Krakencoder’s FCgsr output flavor for the Shen268 parcellation, corresponding to Pearson correlation following global signal regression and matching the preprocessing applied to our oFC data.

### Ridge regression models predicting outcomes

Prediction of post-stroke outcomes in the domains of visual memory, verbal memory, left/right visual attention, language, left/right upper motor, and left/right lower motor were performed using ridge regression with nested cross-validation (CV). Models were built based on connectome information (eSDC, eFC, oFC) in addition to covariates (including age, sex, lesion size, lesion side, days from stroke to scan); for comparison, a covariate only model (COV) that was built using standard linear regression. Models were run separately for each modality, outcome, and timepoint and compared using permutation testing. All connectome models used the Shen268 parcellation atlas, with the upper triangle of each subject’s connectivity matrix vectorized into 35,778 edges. Continuous covariates and connectome edges were z-scored within each training fold and applied to the corresponding test fold to prevent leakage. For a given outcome (e.g. left upper motor), each component score was z-scored within the training set of each fold and averaged into a single composite target per subject.

To ensure comparability across modalities, for a given outcome and timepoint, all four models (eSDC, eFC, oFC, covariate-only) were fit on identical folds/set of subjects. Because outcome scores were missing for varying subsets of subjects across domains and timepoints, sample sizes differed across outcome and timepoint combinations (Supp. Table 1). The outer CV used 5-fold cross-validation repeated 100 times (500 outer folds; 80/20 train-test split in each fold), stratified by quartile bin of the outcome score distribution to ensure balance in train-test splits. Train-test assignments were generated once per outcome/timepoint using a fixed random seed. This ensured that performance differences between modalities could not be attributed to differences in data partitioning. A nested inner 5-fold CV was used for L2 regularization parameter selection across 61 log-spaced values from 10*^−^*^6^ to 10^6^ and optimized using R^2^. Model performances were evaluated using the median Pearson r across outer folds. Statistical significance of model performance was assessed using a permutation test where outcome labels were shuffled and the full nested CV pipeline was rerun 1000 times to generate a null distribution. Model-level permutation p-values were calculated as p = (n + 1)/(N + 1), where n is the number of permuted iterations with median cross-validated r greater than or equal to the observed value and N is the total number of permutations. Resulting p-values were corrected for multiple comparisons using the Benjamini-Hochberg False Discovery Rate^79^ (BH-FDR) procedure. To compare prediction performance between connectome modalities for a given outcome x timepoint we ran permutation-based Wilcoxon rank sum tests on fold level Pearson r values for each of the six pairs among the three connectome modalities and the covariate-only model (5,000 permutations). P-values were corrected using BH-FDR across all tests pooled over timepoints, outcomes and model pairs (q*<*0.05).

Sex-specific models were run identically to the main ridge regression in four configurations: male-train/male-test, female-train/female-test, male-train/female-test, and female-train/male-test. For the cross-sex configurations, males were resampled without replacement to match the female sample size in each repeat, since fewer female participants were available.

### Model feature importances

The Haufe transformation^44^ works by multiplying the beta coefficient vector by the original covariance matrix to reintroduce shared variance among features to the model weights and dividing by the variance of the outcome scores to normalize across outcomes. This allows for more reliable and biologically interpretable model feature importances. ^80^ Here, for each connectome-derived model (eSDC, eFC, oFC) we used the formula *A* = Cov (*X_z_, y*^) */*Var (*y_train_*) – where X_z_ is the z-scored edge feature matrix for a given fold’s training data, ŷ is the model’s predicted output, and Var (*y_train_*) is the variance of the outcome scores among the subjects in that fold’s training set – to compute the Haufe weights per-edge and per fold (35,778 edges over 500 folds for each model, excluding covariates). An edge-level permutation test was performed in each of the 500 outer folds (1000 permutations per fold, same procedure and same runs as the model-level test): training outcome labels were permuted, the ridge model was refit, and the absolute permuted beta weight for each edge was compared with the absolute observed beta weight. The edge-level p-value was calculated as p = (n + 1)/(N + 1), where n is the number of permutations with an absolute permuted beta greater than or equal to the absolute observed beta, and N is the number of permutations. Edges with p*<*0.05 (uncorrected across edges) in at least 400 of the 500 outer folds (80%) were retained to capture the most robust features.^80^ Significant edges’ Haufe values for each outcome and modality pair at the acute timepoint were averaged at the hemispheric-network level, with positive and negative features averaged separately for each hemispheric-network pair and visualized using NiChord.^81^ Network-pair weights in the chord plots were scaled within each modality by the maximum absolute value. Chord width is proportional to the square root of the fraction of edges in the network pair that were retained with that sign, on a common scale across all panels, so that widths are comparable across modalities and outcomes.

### Similarity of feature importance between modalities

To test whether modalities relied on distinct network-level features, we computed the Spearman correlation between the network-level unthresholded (all-fold) Haufe patterns of each pair of modalities (eSDC–eFC, eSDC–oFC, eFC–oFC) for each outcome at the acute time-point. Significance was assessed by permuting the network-pair values of one modality (5,000 permutations) and recomputing *ρ* (two-sided). Given the exploratory nature of this comparison, p-values are reported uncorrected.

All analyses were performed in Python using scikit-learn (version 1.7.2).

## Data availability

The neuroimaging and neuropsychological data used in this study are available at https://cnda.wustl.edu/ap Lesion masks were provided by Dr. Aaron Boes (University of Iowa) and are available upon reasonable request.

## Acknowledgments

This research was supported by NIH RF1 MH123232 (A.K.) and by a grant from the Ann S. Bowers Foundation to C.G. and A.K. through the Ann S. Bowers Women’s Brain Health Initiative.

