## Supplementary Information for "Lesion-Derived Estimated Functional and Structural Connectomes Predict Post-Stroke Recovery Outcomes"

Supplementary Table S1: **Outcome measures by domain and timepoint.** Values are mean ± SD (range). N (acute / 3 months / 1 year) is the number of patients with data for all measures within a domain at that timepoint. Units are given in brackets. Higher scores indicate better performance unless marked ↓ (lower scores indicate better performance).

| Measure | Acute | 3 months | 1 year |
|---|---|---|---|
| **Visual memory** (N = 77 / 68 / 70) | | | |
| BVMT-R immediate recall | $11.88 \pm 8.74$ (0–33) | $16.25 \pm 10.00$ (0–36) | $18.20 \pm 9.10$ (1–36) |
| BVMT-R delayed recall | $4.62 \pm 3.45$ (0–12) | $6.13 \pm 3.95$ (0–12) | $7.03 \pm 3.19$ (1–12) |
| Spatial span forward | $5.58 \pm 2.07$ (2–9) | $5.90 \pm 1.79$ (2–10) | $6.21 \pm 1.74$ (2–10) |
| Spatial span backward | $4.21 \pm 2.40$ (0–9) | $4.90 \pm 2.23$ (0–10) | $5.39 \pm 2.10$ (1–10) |
| **Verbal memory** (N = 70 / 64 / 67) | | | |
| HVLT-R immediate recall | $18.41 \pm 6.10$ (5–31) | $21.05 \pm 5.60$ (10–34) | $21.75 \pm 5.69$ (10–34) |
| HVLT-R delayed recall | $4.51 \pm 3.22$ (0–12) | $6.42 \pm 3.06$ (0–12) | $6.88 \pm 3.28$ (0–12) |
| HVLT-R discrimination index | $8.16 \pm 2.81$ (0–12) | $9.14 \pm 2.27$ (2–12) | $9.69 \pm 2.20$ (3–12) |
| **Left attention** (N = 79 / 71 / 66) | | | |
| Posner accuracy, left invalid [proportion] | $0.91 \pm 0.17$ (0.1–1.0) | $0.97 \pm 0.08$ (0.5–1.0) | $0.96 \pm 0.13$ (0.1–1.0) |
| Posner accuracy, left valid [proportion] | $0.93 \pm 0.13$ (0.1–1.0) | $0.96 \pm 0.07$ (0.7–1.0) | $0.96 \pm 0.13$ (0.0–1.0) |
| Posner RT, left invalid [ms] ↓ | $622 \pm 224$ (299–1364) | $584 \pm 177$ (286–1495) | $591 \pm 192$ (282–1324) |
| Posner RT, left valid [ms] ↓ | $588 \pm 219$ (302–1206) | $556 \pm 159$ (310–1218) | $565 \pm 167$ (281–1297) |
| Mesulam cancellation, left omissions [count] ↓ | $3.73 \pm 7.02$ (0–30) | $1.06 \pm 1.79$ (0–9) | $1.02 \pm 2.18$ (0–14) |
| BIT star cancellation, left omissions [count] ↓ | $1.03 \pm 2.34$ (0–13) | $0.56 \pm 1.83$ (0–14) | $0.29 \pm 0.72$ (0–4) |
| **Right attention** (N = 79 / 71 / 66) | | | |
| Posner accuracy, right invalid [proportion] | $0.94 \pm 0.13$ (0.2–1.0) | $0.97 \pm 0.08$ (0.6–1.0) | $0.98 \pm 0.06$ (0.6–1.0) |
| Posner accuracy, right valid [proportion] | $0.95 \pm 0.12$ (0.1–1.0) | $0.97 \pm 0.08$ (0.4–1.0) | $0.98 \pm 0.03$ (0.8–1.0) |
| Posner RT, right invalid [ms] ↓ | $623 \pm 261$ (344–2000) | $570 \pm 178$ (312–1336) | $557 \pm 171$ (293–1401) |
| Posner RT, right valid [ms] ↓ | $568 \pm 197$ (318–1300) | $558 \pm 180$ (304–1443) | $544 \pm 150$ (269–1035) |
| Mesulam cancellation, right omissions [count] ↓ | $2.61 \pm 5.29$ (0–30) | $0.94 \pm 1.66$ (0–8) | $0.92 \pm 1.86$ (0–13) |
| BIT star cancellation, right omissions [count] ↓ | $0.95 \pm 2.14$ (0–14) | $0.35 \pm 0.76$ (0–3) | $0.32 \pm 0.77$ (0–4) |
| **Language** (N = 92 / 74 / 68) | | | |
| Word reading | $34.61 \pm 4.57$ (11–37) | $35.72 \pm 2.00$ (22–37) | $35.91 \pm 1.22$ (31–37) |
| Commands | $13.59 \pm 3.16$ (0–15) | $14.39 \pm 2.05$ (0–15) | $14.66 \pm 0.92$ (9–15) |
| Boston Naming Test | $12.37 \pm 3.52$ (0–15) | $13.27 \pm 2.41$ (0–15) | $13.57 \pm 1.87$ (6–15) |
| Complex ideational material | $8.92 \pm 3.26$ (0–12) | $10.16 \pm 2.34$ (0–12) | $10.65 \pm 1.82$ (3–12) |
| Reading | $7.98 \pm 3.19$ (0–10) | $8.99 \pm 1.96$ (0–10) | $9.31 \pm 1.31$ (3–10) |
| Reading comprehension | $3.89 \pm 1.57$ (0–5) | $4.57 \pm 0.94$ (0–5) | $4.62 \pm 0.88$ (0–5) |

Supplementary Table S1: **Outcome measures by domain and timepoint (continued).**

| Measure | Acute | 3 months | 1 year |
|---|---|---|---|
| **Left upper motor** (N = 94 / 75 / 67) | | | |
| Shoulder flexion AROM, L [deg] | $143.4 \pm 60.3$ (0–180) | $152.7 \pm 48.8$ (0–180) | $157.2 \pm 37.1$ (0–180) |
| Wrist extension AROM, L [deg] | $52.2 \pm 24.2$ (0–90) | $53.2 \pm 21.8$ (0–90) | $49.3 \pm 19.1$ (0–80) |
| Grip strength, L [kg] | $21.2 \pm 14.5$ (0–60) | $20.0 \pm 13.8$ (0–53) | $21.8 \pm 12.8$ (0–45) |
| 9-hole peg test, L [pegs/s] | $0.25 \pm 0.15$ (0.0–0.5) | $0.28 \pm 0.15$ (0.0–0.5) | $0.29 \pm 0.14$ (0.0–0.6) |
| Action Research Arm Test, L | $46.0 \pm 20.3$ (0–57) | $48.7 \pm 17.0$ (0–57) | $50.9 \pm 14.8$ (0–57) |
| **Right upper motor** (N = 94 / 75 / 67) | | | |
| Shoulder flexion AROM, R [deg] | $149.8 \pm 57.0$ (0–180) | $161.6 \pm 39.2$ (0–180) | $162.5 \pm 37.9$ (0–180) |
| Wrist extension AROM, R [deg] | $50.2 \pm 23.7$ (0–90) | $57.3 \pm 19.1$ (0–85) | $51.4 \pm 18.0$ (0–90) |
| Grip strength, R [kg] | $22.3 \pm 15.1$ (0–69) | $23.0 \pm 12.7$ (0–68) | $23.3 \pm 12.1$ (0–45) |
| 9-hole peg test, R [pegs/s] | $0.27 \pm 0.16$ (0.0–0.5) | $0.33 \pm 0.12$ (0.0–0.6) | $0.34 \pm 0.12$ (0.0–0.5) |
| Action Research Arm Test, R | $46.8 \pm 19.6$ (0–57) | $52.9 \pm 13.2$ (0–57) | $53.2 \pm 13.3$ (0–57) |
| **Left lower motor** (N = 73 / 66 / 65) | | | |
| Ankle dorsiflexion AROM, L [deg][a] | $12.6 \pm 6.3$ (0–25) | $13.1 \pm 5.7$ (0–30) | $12.1 \pm 5.2$ (0–22) |
| Knee extension AROM, L [deg][b] | $-6.6 \pm 22.8$ (−90–0) | $-4.1 \pm 16.1$ (−90–0) | $-4.4 \pm 18.4$ (−90–0) |
| Hip flexion AROM, L [deg][c] | $112.5 \pm 34.7$ (0–165) | $112.7 \pm 28.4$ (0–160) | $109.9 \pm 23.8$ (0–150) |
| Ankle strength, L | $28.6 \pm 10.0$ (0–33) | $30.2 \pm 8.2$ (0–33) | $30.2 \pm 7.8$ (0–33) |
| Knee strength, L | $29.2 \pm 9.5$ (0–33) | $30.8 \pm 6.3$ (0–33) | $30.9 \pm 6.4$ (0–33) |
| Hip strength, L | $27.5 \pm 10.0$ (0–33) | $29.4 \pm 7.3$ (0–33) | $29.5 \pm 6.9$ (0–33) |
| **Right lower motor** (N = 73 / 66 / 65) | | | |
| Ankle dorsiflexion AROM, R [deg][a] | $12.8 \pm 5.9$ (0–25) | $13.2 \pm 6.4$ (0–40) | $12.9 \pm 4.6$ (0–20) |
| Knee extension AROM, R [deg][b] | $-5.8 \pm 20.0$ (−90–0) | $-2.2 \pm 10.6$ (−80–0) | $-1.6 \pm 7.7$ (−58–0) |
| Hip flexion AROM, R [deg][c] | $114.4 \pm 32.0$ (0–160) | $117.4 \pm 20.5$ (0–160) | $116.3 \pm 13.5$ (95–160) |
| Ankle strength, R | $29.0 \pm 9.0$ (0–33) | $31.6 \pm 6.0$ (0–33) | $30.3 \pm 8.3$ (0–33) |
| Knee strength, R | $29.7 \pm 8.4$ (0–33) | $32.4 \pm 2.4$ (19–33) | $32.1 \pm 3.0$ (19–33) |
| Hip strength, R | $28.3 \pm 8.8$ (0–33) | $31.2 \pm 5.0$ (0–33) | $30.1 \pm 4.7$ (14–33) |

AROM, active range of motion. [a]Typical range in healthy adults approximately 5–20°. [b]0° indicates full knee extension (typical in healthy adults); negative values indicate incomplete extension. [c]Typical range in healthy adults approximately 100–150°.

Supplementary Table S2: **Sample sizes for the sex-specific and cross-sex analyses.** Participants with complete data for each outcome and timepoint. In each of 100 repeats, males were randomly subsampled to the number of females, and 5-fold cross-validation was performed within each sex, so training and test set sizes were identical for male and female models. Training and test $n$ are per sex per fold; ranges reflect folds of unequal size when $n$ was not divisible by 5. Left and right outcomes used identical participants.

| Outcome | Timepoint | Males | Females | Training $n$ | Test $n$ |
|---|---|---|---|---|---|
| Visual memory | Acute | 43 | 34 | 27–28 | 6–7 |
| | 3 mo | 39 | 29 | 23–24 | 5–6 |
| | 1 yr | 39 | 31 | 24–25 | 6–7 |
| Verbal memory | Acute | 41 | 29 | 23–24 | 5–6 |
| | 3 mo | 37 | 27 | 21–22 | 5–6 |
| | 1 yr | 38 | 29 | 23–24 | 5–6 |
| Attention (left, right) | Acute | 43 | 36 | 28–29 | 7–8 |
| | 3 mo | 37 | 34 | 27–28 | 6–7 |
| | 1 yr | 35 | 31 | 24–25 | 6–7 |
| Language | Acute | 52 | 40 | 32 | 8 |
| | 3 mo | 41 | 33 | 26–27 | 6–7 |
| | 1 yr | 38 | 30 | 24 | 6 |
| Upper motor (left, right) | Acute | 54 | 40 | 32 | 8 |
| | 3 mo | 40 | 35 | 28 | 7 |
| | 1 yr | 38 | 29 | 23–24 | 5–6 |
| Lower motor (left, right) | Acute | 42 | 31 | 24–25 | 6–7 |
| | 3 mo | 37 | 29 | 23–24 | 5–6 |
| | 1 yr | 37 | 28 | 22–23 | 5–6 |

Supplementary Table S3: **Stability of network-level feature importance across timepoints.** Spearman $\rho$ between network-pair-averaged Haufe patterns of the same modality and outcome at two timepoints. Values are descriptive because outcomes at all timepoints were predicted from the same baseline imaging metric (lesion-derived eSDC and eFC, and acute observed FC). A dash indicates the model was not significant against its permutation null (BH-FDR $q < 0.05$) at one or both timepoints.

| Modality | Outcome | Acute vs 3 mo | Acute vs 1 yr | 3 mo vs 1 yr |
|---|---|---|---|---|
| eSDC | Visual memory | – | – | – |
| | Verbal memory | – | – | – |
| | Attention (L) | 0.960 | – | – |
| | Attention (R) | 0.923 | – | – |
| | Language | 0.978 | 0.914 | 0.944 |
| | Upper motor (L) | 0.993 | 0.991 | 0.998 |
| | Upper motor (R) | 0.996 | 0.986 | 0.994 |
| | Lower motor (L) | 0.992 | 0.994 | 0.999 |
| | Lower motor (R) | 0.991 | 0.978 | 0.992 |
| eFC | Visual memory | – | – | – |
| | Verbal memory | – | – | – |
| | Attention (L) | 0.983 | – | – |
| | Attention (R) | 0.985 | – | – |
| | Language | 0.955 | 0.908 | 0.982 |
| | Upper motor (L) | 0.974 | 0.988 | 0.988 |
| | Upper motor (R) | 0.963 | 0.973 | 0.996 |
| | Lower motor (L) | 0.986 | 0.994 | 0.993 |
| | Lower motor (R) | 0.988 | 0.959 | 0.968 |
| oFC | Visual memory | 0.450 | 0.389 | 0.737 |
| | Verbal memory | – | – | – |
| | Attention (L) | – | – | – |
| | Attention (R) | – | – | – |
| | Language | 0.676 | 0.743 | 0.801 |
| | Upper motor (L) | 0.928 | 0.928 | 0.945 |
| | Upper motor (R) | 0.796 | – | – |
| | Lower motor (L) | 0.918 | 0.915 | 0.978 |
| | Lower motor (R) | – | – | – |

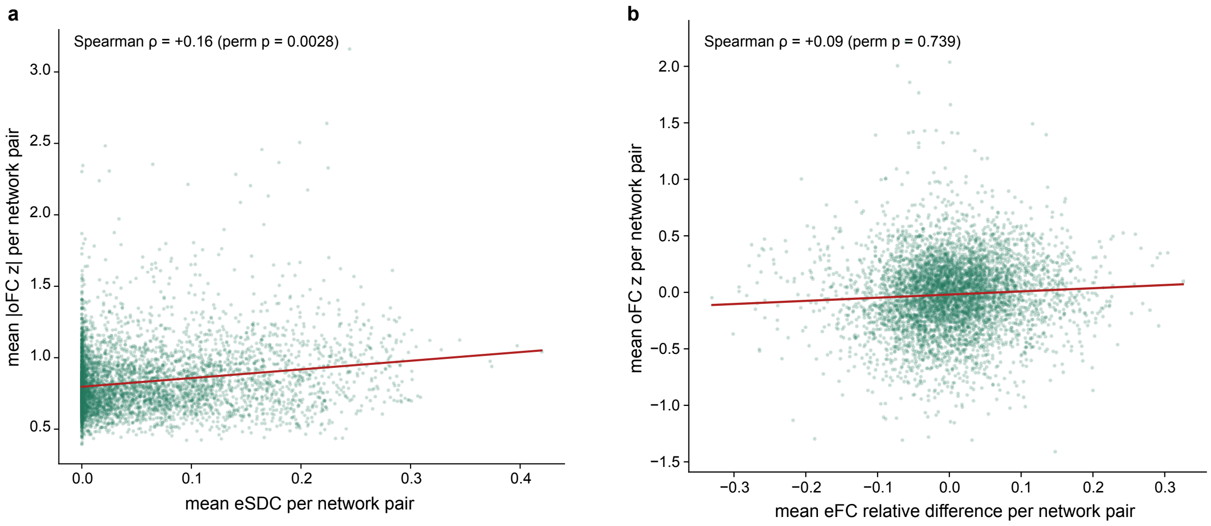


Supplementary Figure S1: **Individual-level correlation between network-pair disruption.** Each point is one network pair for one stroke patient (acute timepoint; 96 patients $\times$ 55 network pairs = 5,280 points per panel). Network pairs are all unique pairs of 10 networks, including self-pairs: the Yeo 7 networks plus subcortical, cerebellar and brainstem regions. Values are averaged over the edges within each network pair for each stroke patient. **a)** Mean absolute oFC $z$-score (relative to 22 healthy controls) versus mean eSDC (percent disruption). **b)** Mean oFC $z$-score versus mean eFC relative difference from the HCP-YA reference (N=420).

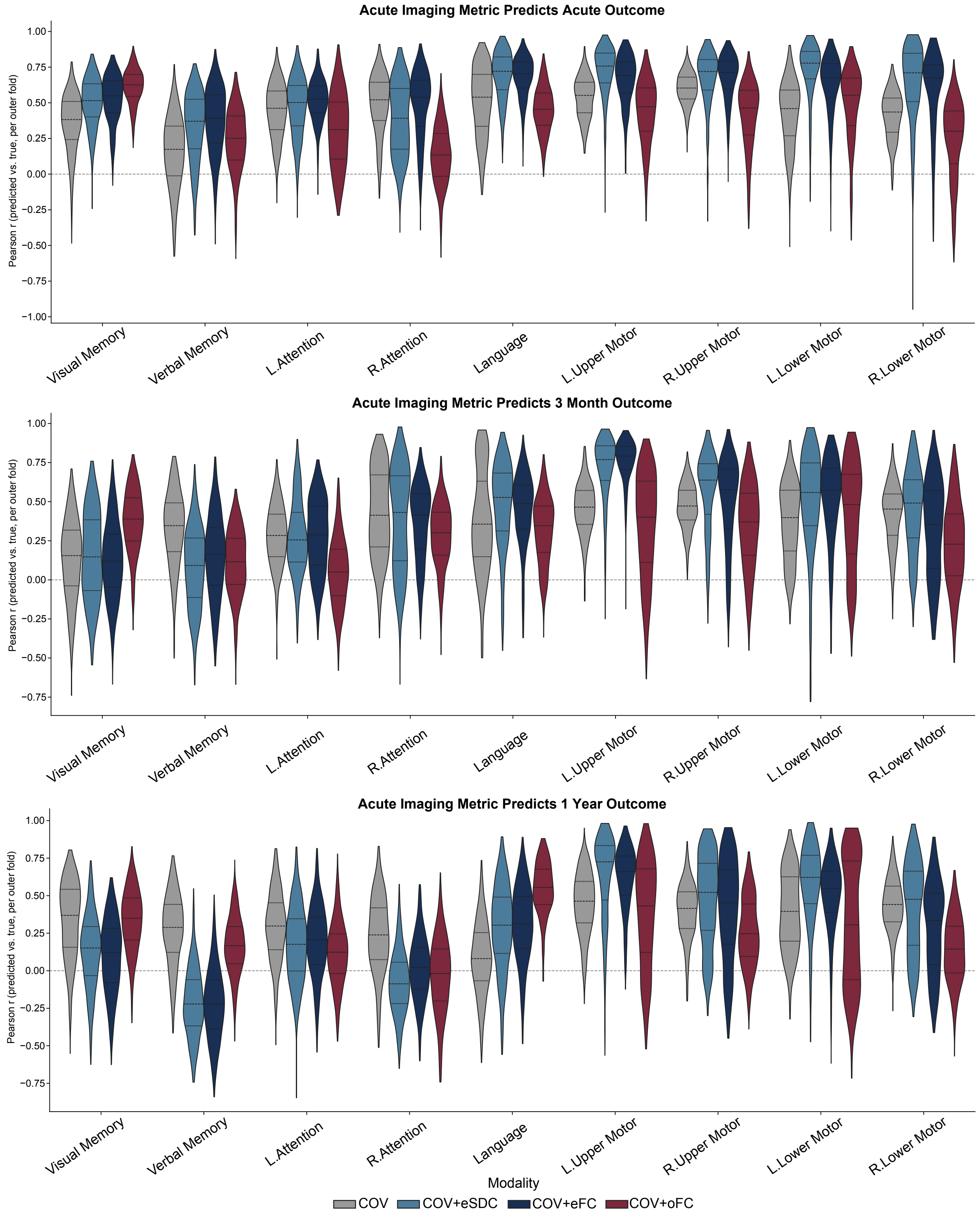


Supplementary Figure S2: **Fold-level prediction performance for all outcomes and timepoints.** Violins show Pearson $r$ across 5-fold cross-validation repeated 100 times (500 outer folds) for the covariate-only, eSDC, eFC and oFC ridge regression model predictions at each timepoint.

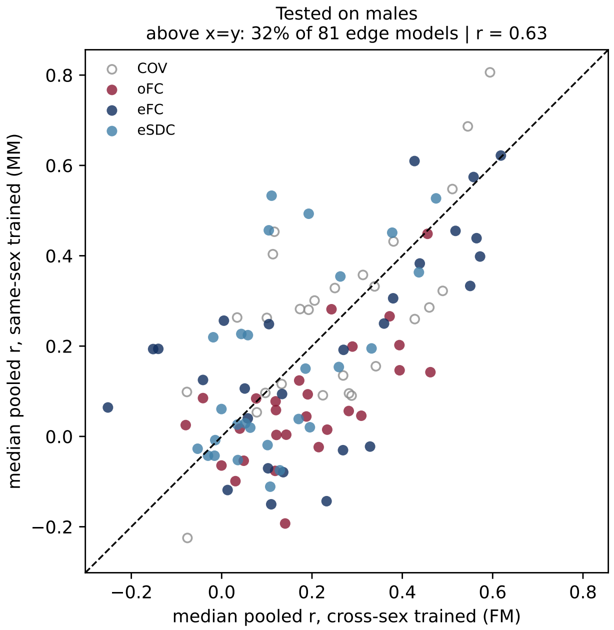


Supplementary Figure S3: **Same-sex versus cross-sex training.** Models were trained on male-only or female-only data and tested on the same held-out participants, with males randomly subsampled to the female sample size in each of 100 repeats of 5-fold cross-validation, so training and test set sizes were identical across sexes. Each point is one outcome × modality × timepoint (median pooled Pearson r across repeats). Left: tested on males (y, male-trained; x, female-trained). Right: tested on females (y, female-trained; x, male-trained). Points above the line indicate better prediction after same-sex training. Because both models in a panel are evaluated on identical test participants, differences between axes reflect training sex rather than differences in test-set score distributions; comparisons should be made within, not across, panels. Gray open circles indicate covariate-only models.

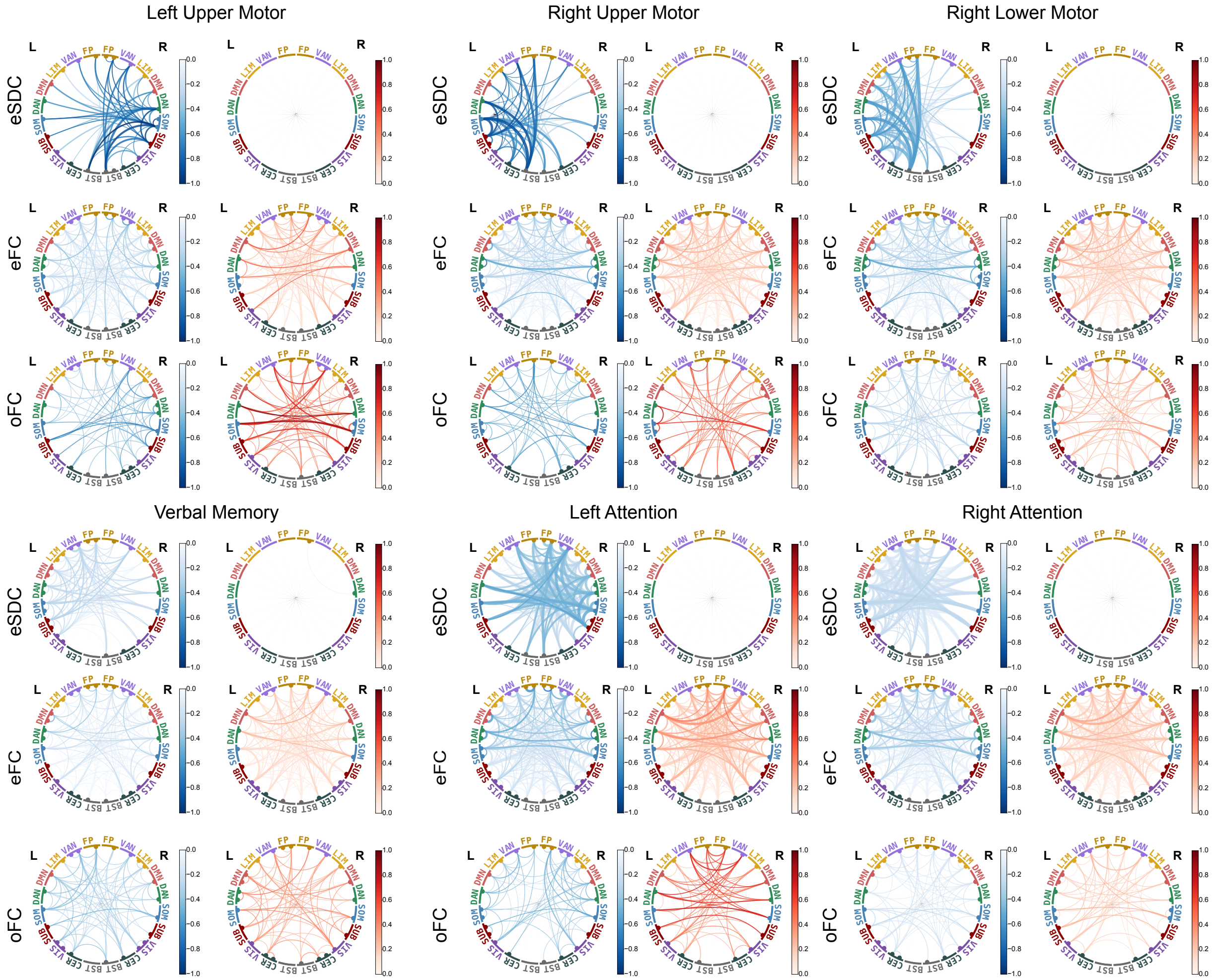


Supplementary Figure S4: **Network-pair feature importances for models predicting acute post-stroke outcomes.** Network-level feature importances driving predictions for left/right upper motor, right lower motor, verbal memory, and left/right attention outcomes at the acute timepoint, shown separately for eSDC, eFC, and oFC models (top, middle and bottom, respectively). Chord plots show network-pair averages of significant positive predictive features (hot colorbar) where larger (dis)connectome values indicate better behavioral scores and significant negative predictive features (cool colorbar) where larger (dis)connectome values indicate worse behavioral scores. Chord width is proportional to the square root of the fraction of edges in each network pair that met the stability criterion with that sign, on a common scale across all panels. L and R denote left and right hemispheric networks.
